# Genetic Studies Highlight the Role of TET2 and INO80 in DNA Damage Response and Kidney Disease Pathogenesis

**DOI:** 10.1101/2024.02.02.578718

**Authors:** Xiujie Liang, Hongbo Liu, Hailong Hu, Jianfu Zhou, Amin Abedini, Andrea Sanchez Navarro, Konstantin A. Klötzer, Katalin Susztak

## Abstract

Genome-wide association studies (GWAS) have identified over 800 loci associated with kidney function, yet the specific genes, variants, and pathways involved remain elusive. By integrating kidney function GWAS, human kidney expression and methylation quantitative trait analyses, we identified Ten-Eleven Translocation (TET) DNA demethylase 2: TET2 as a novel kidney disease risk gene. Utilizing single-cell chromatin accessibility and CRISPR-based genome editing, we highlight GWAS variants that influence *TET2* expression in kidney proximal tubule cells.

Experiments using kidney-tubule-specific *Tet2* knockout mice indicated its protective role in cisplatin-induced acute kidney injury, as well as chronic kidney disease and fibrosis, induced by unilateral ureteral obstruction or adenine diet. Single-cell gene profiling of kidneys from *Tet2* knockout mice and *TET2-*knock-down tubule cells revealed the altered expression of DNA damage repair and chromosome segregation genes, notably including *INO80*, another kidney function GWAS target gene itself.

Remarkably both *TET2-*null and *INO80-*null cells exhibited an increased accumulation of micronuclei after injury, leading to the activation of cytosolic nucleotide sensor cGAS-STING. Genetic deletion of cGAS or STING in kidney tubules or pharmacological inhibition of STING protected TET2 null mice from disease development. In conclusion, our findings highlight TET2 and INO80 as key genes in the pathogenesis of kidney diseases, indicating the importance of DNA damage repair mechanisms.

## Introduction

Kidney disease is a major global health burden, affecting over 850 million people worldwide^1^. Kidney disease incidence is much higher in older adults and it is one of the fastest-growing causes of mortality^2^. A better understanding of the mechanism of kidney dysfunction is crucial for developing new therapeutics to treat or cure kidney disease.

Kidney function shows strong heritability^3–5^. Large genome-wide association studies (GWAS) of kidney function have identified more than 800 loci that demonstrate statistically significant and reproducible association with kidney function ^6, 7^. However, over 90% of variants identified by GWAS reside in the noncoding regions of the genome^8^, which are often correlated due to strong linkage disequilibrium (LD)^9^. Consequently, pinpointing the causal variants, their target genes, specific cell types involved, and underlying disease mechanisms continues to be a major challenge.

Disease-causing variants are often located in cell type-specific regulatory regions, potentially altering the binding strength of transcription factors and resulting in quantitative differences in the expression of cell type-specific target genes. The expression quantitative trait loci (eQTL) analysis, which defines the association between genetic variants and tissue gene expression, has been widely used to identify target genes in GWAS^3, 10–12^. Analyzing the effects of genotype on the epigenome, such as DNA methylation, can improve gene and cell type prioritization^7^. In addition, examining changes in gene expression and regulation at a single-cell level, rather than at the whole tissue bulk level, is critical. While computational prioritization strategies have prioritized new kidney function risk genes based on GWAS and human kidney multi-omics datasets, these studies need to be followed by careful cell and animal model experiments to substantiate the causal role of a specific gene.

DNA methylation, the addition of a methyl group to the 5-position of cytosines, is mediated by DNA methyltransferases (DNMT). The Ten-Eleven Translocation (TET) enzymes play a role in demethylation by converting 5-methylcytosine (5mC) to 5-hydroxymethylcytosine (5hmC), 5-formylcytosine, and 5-carboxylcytosine^13, 14^. DNA methylation is an energetically costly process which also increase the mutation rate as 5mC is converted to thymine, which is difficult to repair. DNA methylation is an important regulator of multiple DNA-based processes. Most studies have analyzed the role of cytosine methylation in gene expression regulation. Cytosine methylation renders the DNA region less accessible to the transcriptional machinery by recruiting methyl binding proteins and altering transcription factor binding strength^15^. Cytosine methylation is critical during kidney development and our previous studies demonstrated the key role of *Dnmt1* in Six2 in progenitor cells as it is required for transposable element silencing ^16^ while *Dnmt3a* and *Dnmt3b* are essential for the silencing developmental genes^17^. Kidneys obtained from patients with CKD and fibrosis also showed important changes in cytosine methylation at more than 100 CpG sites ^18^. Globally more CpG sites show higher methylation level in CKD. These methylation changes correlate and predict kidney function decline indicating their potential role in disease development.

DNA damage occurs in various forms, including abasic sites, adducts, DNA-protein cross-links, insertion/deletion mismatches, double-strand breaks (DSBs), and single-strand breaks (SSBs) ^19^. Cells respond to DNA damage by initiating a series of highly coordinated events to repair DNA, known as the DNA damage response (DDR) pathway, which includes base excision repair (BER), nucleotide excision repair (NER), mismatch repair (MMR), non-homologous end joining (NHEJ), and homologous recombination (HR) ^20^. The ataxia teleangiectasia mutated (ATM) and ATM-RAD3 related (ATR) and the DNA-dependent protein (DNA-PK) are three kinases that control DDR and orchestrate DSB. In cases where the DNA damage exceeds the repair capacity of the cell, the DDR triggers senescence or apoptosis. By arresting proliferation and promoting clearance of damaged cells, the DDR acts as barrier to tumorigenesis. Recent studies highlighted the role of ATM and ATR in kidney disease development. Mutations in DNA repair pathway genes, such as *ERCC1*, *ERCC2*, *ERCC6,* and *ERCC8*, have been associated with progeria, cancer, or immunodeficiency^21^. The modified *ERCC1* knockout mouse model, which expresses a liver-specific *ERCC1* rescue transgene, developed proteinuria, segmental glomerulosclerosis, and end-stage renal failure ^21, 22^. The role of cytosine methylation in DDR is not fully understood, however both epigenetic changes and DNA damage is strongly associated with aging.

In this study, we identified genetic variants associated with kidney function that concurrently lower the expression of *Tet2* in kidney proximal tubule cells. Kidney-specific gene knockout mice have highlighted the protective roles of *Tet2* both in acute kidney injury and chronic fibrosis. Molecular studies indicated the role of *TET2* in the HR pathway for DSB repair and in ensuring proper chromosome segregation in proximal tubule cells. Loss of *Tet2* lead to kidney injury by leaving micronuclei in the cytosol, resulting in cGAS-STING activation and subsequent kidney disease development.

## Results

### Prioritization of TET2 for kidney function

Multiple GWAS have identified common noncoding nucleotide variants on chromosome 4 associated with kidney function^4, 23–25,7^. To prioritize the causal variants within this locus, we performed statistical fine mapping using the SuSie method. We identified two credible sets: rs6533181 (posterior inclusion probability, PIP = 9.98 × 10^-8^), rs115918101 (PIP = 8.82 × 10^-8^), and rs2214407 (PIP = 3.54 × 10^-8^) (Figure 1A). The variant rs6533181 was significantly associated with kidney function in the eGFR creatinine GWAS (*P* =3.83 × 10^-11^) (Figure 1B) ^7^.

**Figure 1.**
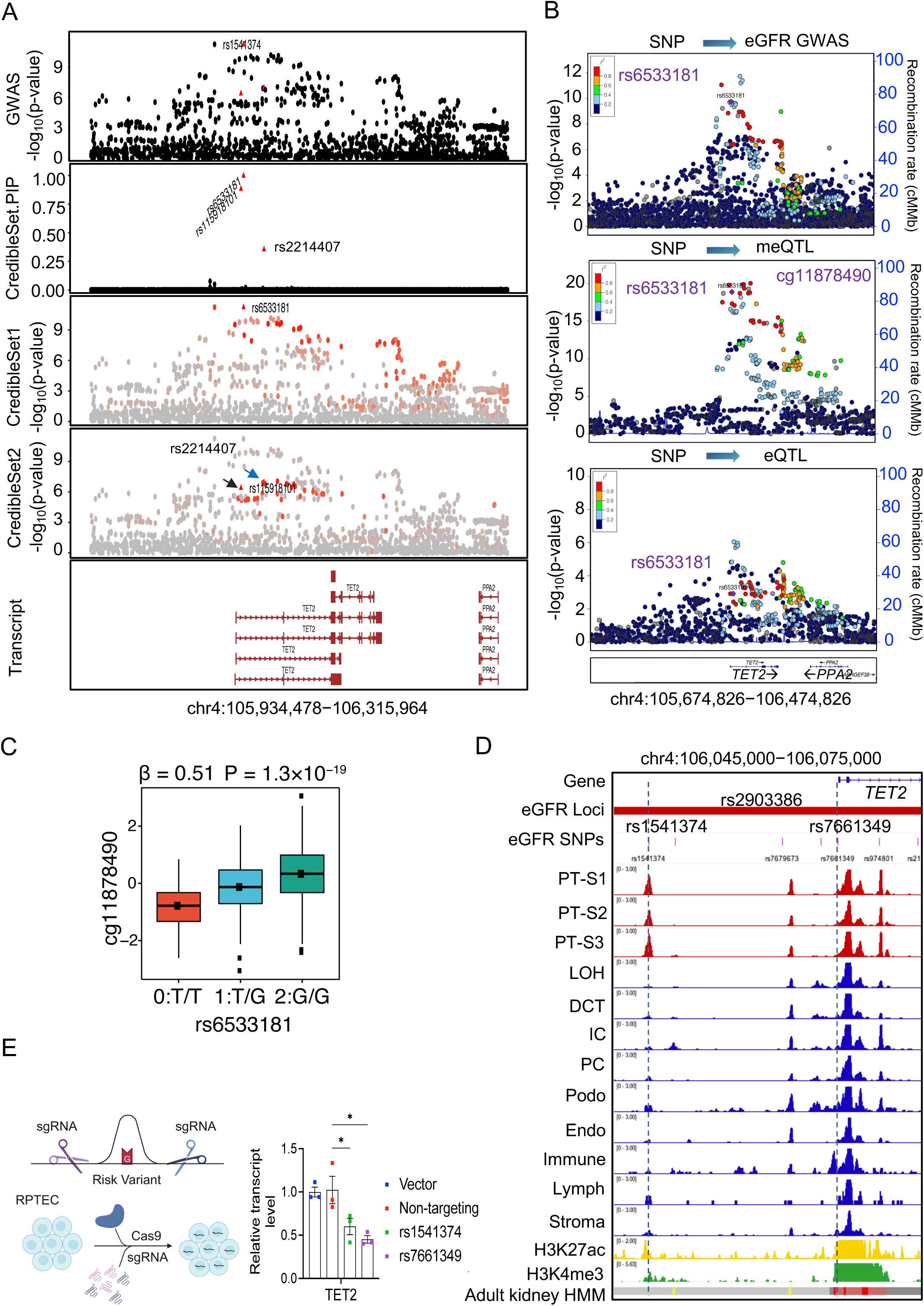
Identification of *TET2* as a kidney disease risk gene. **(A)** LocusZoom plots of eGFR GWAS (genotype and eGFRcrea association, n = 1,746,932) and fine mapping plots for the region chr4: 105,934,478-106,315,964. Each dot in the plot represents a single SNP (Single Nucleotide Polymorphism). The color indicates the LD association. **(B)** LocusZoom plots of GWAS (genotype and eGFRcrea association, n = 1,508,659), kidney CpG cg11878490 meQTL (genotype and cg11878490 methylation association, n = 443), and kidney *TET2* eQTLs (genotype and *TET2* expression association, n = 686). The y-axis displays the -log_10_(*p*-values) of association tests from GWAS, meQTL, and eQTLs studies. **(C)** Genotype (rs6533181, x-axis) and normalized CpG methylation (cg11878490, y-axis) association in human kidneys. Center lines show the medians, and the box limits indicate the 25^th^ and 75^th^ percentiles. *P*-value was calculated using a linear regression meQTL model. **(D)** From top to bottom: Gene browser view of the eGFR GWAS SNPs within the specified regions; single-nucleus Assay for Transposase-Accessible Chromatin using sequencing (snATAC-seq) analysis of chromatin accessibility of human kidneys. S1 segment of proximal tubules (PT-S1), S2 segment of proximal tubules (PT-S2), S3 segment of proximal tubules (PT-S3), loop of Henle (LOH), distal convoluted tubule (DCT), connecting tubule (CNT), intercalated cells (IC), collecting duct principal cell types (PC), podocytes (Podo), endothelial cells (Endo), immune cells (Immune), lymphocyte (Lymph); human kidney histone modifications by chromatin immunoprecipitation (ChIP)-seq and human adult kidney chromatin states. **(E)** Relative transcript levels of *TET2* after CRISPR-mediated deletion of the locus, with a sample size of n = 3. *GAPDH* was used for normalization. \**p* < 0.05 versus indicated group.

To further dissect this region, we integrated the GWAS data with kidney methylation quantitative trait loci (meQTLs) and allele-specific expression (ASE) analyses. We observed a significant association between the variant rs6533181 and CpG methylation (cg11878490 meQTLs, *P* = 1.29 × 10^-19^) and *TET2* expression in kidney tubule samples (tubule ASE, *P* = 9.66 × 10^-5^) (Figure 1, B and C, and Supplemental Figure 1A). Multi-tissue eQTL analysis from the GTEx project (Genotype-Tissue Expression) indicated that rs6533181 is associated with *TET2* expression levels in the skin (*P* =1.2 × 10^-10^) and in fibroblasts (*P* =6.1 × 10^-5^).

To test whether eGFR, cytosine methylation of cg11878490, and *TET2* expression share causal variants at this locus, we performed a statistical colocalization analysis using Moloc. We found compelling evidence that the variants linked with kidney function, CpG methylation, and *TET2* expression in the kidney were shared, as indicated by a high posterior probability (ASE.PP.H4.abf = 0.87). *TET2* expression in kidney tubules was strongly genotype-dependent (rs6533181, eGFR GWAS risk allele T, ASE *P* = 9.66 × 10^-5^) (Supplemental Figure 1A). To further identify target genes for this variant, we integrated evidence from activity-by-contact (ABC) model ^26^. The score from the ABC model for this variant was 0.4468. According to the combined SNP-to-gene (cS2G) linking strategy^27^, the association score for this variant and *TET2* was 1, strongly suggesting that *TET2* is the target gene of this genetic locus.

Considering that SNPs located in the regulatory region are more likely to be causal^28, 29^, we employed our human kidney single-nuclear assay for transposase-accessible chromatin sequencing (snATAC-seq) to further prioritize risk variants. The variant rs6533181 was not located in an open chromatin region in the human kidney (Supplemental Figure 1B), however, rs1541374 and rs7661349, which are in very strong LD with rs6533181 (r >0.93 and r > 0.98, respectively), were located within an open chromatin region in the proximal tubule cell and multiple cell types, respectively (Figure 1D). These observations prompted us to hypothesize that rs7661349 and rs1541374 are the likely causal variants. To demonstrate that rs7661349 and rs1541374 regulate the expression of *TET2* in human kidney proximal tubule cells, we performed CRISPR-based deletion of open chromatin regions containing these two variants (Figure 1E and Supplemental Figure 2A). Deletion of rs7661349 and rs1541374, but not of the non-target regions, lowered *TET2* levels, but not *Inorganic pyrophosphatase 2* (*PPA2*) levels (Figure 1E and Supplemental Figure 2B).

Our human kidney single-nuclear RNA sequencing (snRNA-seq) data showed that *TET2* expression in the proximal tubule (S3 segment) was lower in diseased kidneys compared to healthy kidneys (Supplemental Figure 3A). Immunofluorescence staining further confirmed that the protein level of TET2 in the proximal tubule was lower in diseased kidneys compared to controls (Supplemental Figure 3C).

In summary, by integrating genetic analysis, single-nucleus epigenome mapping, and CRISPR-Cas9 gene editing, we identified *TET2* as a risk gene for kidney disease, specifically within kidney proximal tubule cells, where lower *TET2* expression was associated with an elevated risk of renal disease.

### Kidney-specific Tet2 knockout mice show increased severity of acute and chronic kidney disease

To study the role of *TET2* in kidney disease development, we generated mice with genetically lowered *Tet2* expression in kidney cells by breeding *Six2^Cre^* mice with *Tet2^f/f^* mice^30^. Gene expression analysis confirmed the reduction in *Tet2* expression in the proximal tubules of the kidneys in adult *Six2^Cre^Tet2^f/f^* mice compared to *Six2^Cr^*^e^ mice (Supplemental Figure 4, A and B). *Six2^Cre^Tet2^f/f^* mice were born at the expected Mendelian ratio, appeared healthy at birth, and showed no significant differences in life span at 40 weeks of age ^30^. Next, we analyzed *Six2^Cre^Tet2^wt/f^* and *Six2^Cre^Tet2^f/f^* mice in a cisplatin-induced acute kidney injury (AKI) model (Figure 2A). Blood urea nitrogen (BUN) levels were elevated in cisplatin-injected *Six2^Cre^Tet2^wt/f^* and *Six2^Cre^Tet2^f/f^* mice compared to cisplatin-injected *Six2^Cre^* mice (Figure 2B). Histological analysis indicated more hyaline casts, loss of bush border, and tubular lumen dilation in kidneys of cisplatin-injected *Six2^Cre^Tet2^wt/f^* and *Six2^Cre^Tet2^f/f^* mice compared to those in cisplatin-injected control mice (Figure 2C). Expression levels of kidney injury markers, such as *kidney injury molecule* 1 (*Kim1*) and *neutrophil gelatinase-associated lipocalin* (*Ngal*), were higher in kidneys of cisplatin-injected *Six2^Cre^Tet2^wt/f^* and *Six2^Cre^Tet2^f/f^* mice when compared with cisplatin-injected control *Six2^Cre^* mice (Figure 2D).

**Figure 2.**
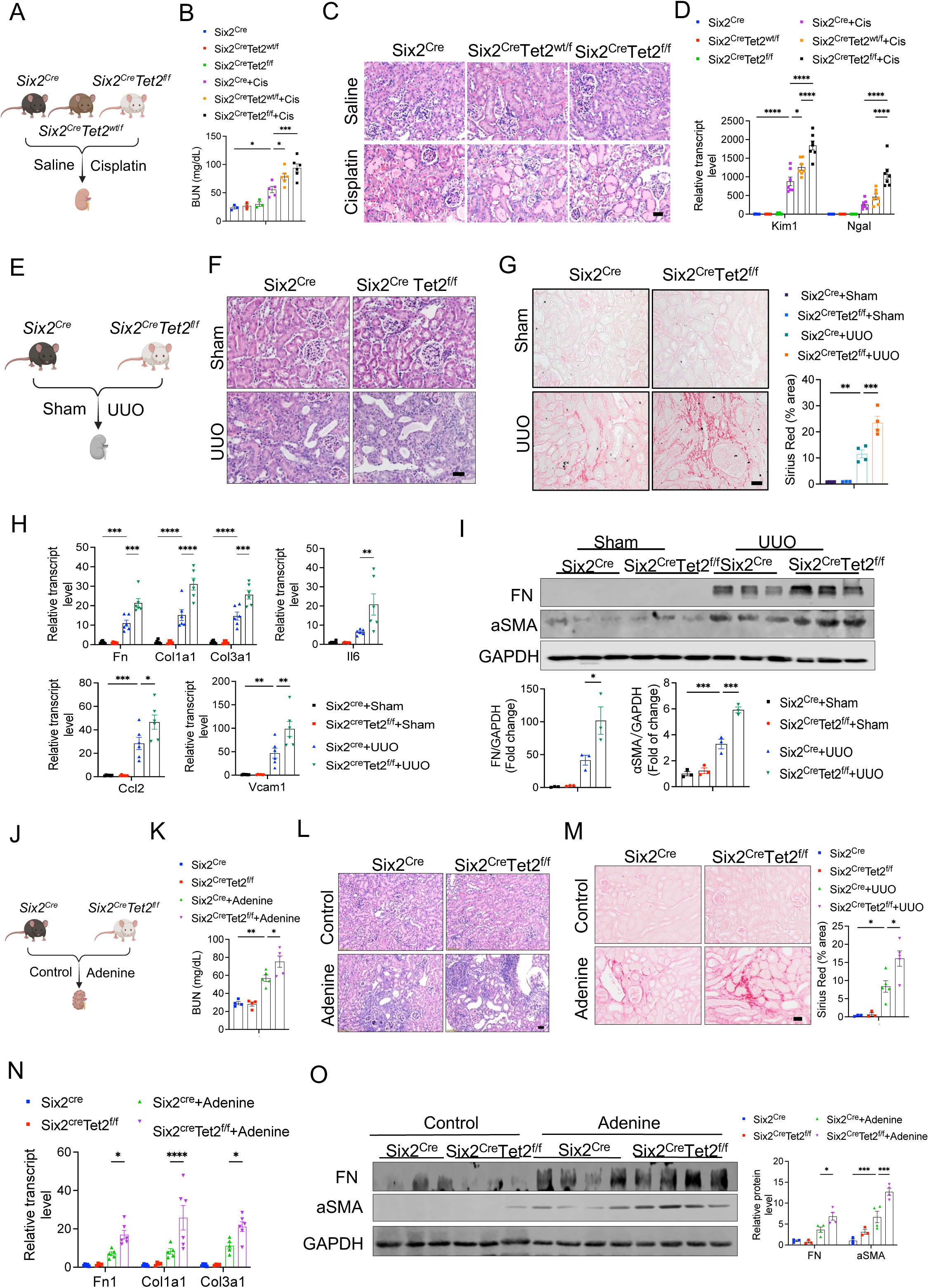
Tubule-specific *Tet2* loss exacerbated renal injury and fibrosis. **(A)** Experimental design. *Six2^Cre^*, *Six2^cre^Tet2^wt/f^*, and *Six2^cre^Tet2^f/f^* mice were injected with saline or cisplatin. Mice were euthanized 3 days after injection. **(B)** BUN levels were measured in *Six2^Cre^*, *Six2^cre^Tet^wt/f^*, and *Six2^cre^Tet2^f/f^* mice injected with saline or cisplatin. In the saline-injected group: n = 3. In the cisplatin-injected group: *Six2^Cre^* (n = 5), *Six2^cre^Tet^wt/f^* (n = 6), and *Six2^cre^Tet2^f/f^* (n = 6). **(C)** Representative images of H&E-stained kidneys sections from *Six2^Cre^* and *Six2^cre^Tet2^f/f^* mice injected with saline or cisplatin. **(D)** Relative transcript level of *Kim1* and *Ngal* in kidneys of *Six2^Cre^* and *Six2^cre^Tet2^f/f^* mice injected with saline or cisplatin. *Gapdh* was used for normalization. In the saline-injected group: *Six2^Cre^* (n = 6), *Six2^cre^Tet^wt/f^* (n = 4), and *Six2^cre^Tet2^f/f^* (n = 6). In the cisplatin-injected groups: n = 5 each. **(E)** Experimental setup. *Six2^Cre^* and *Six2^cre^Tet2^f/^*^f^ mice were subjected to sham-operation or unilateral ureteral obstruction (UUO). Mice were euthanized 4 days after surgery. **(F)** Representative images of H&E-stained kidney sections from *Six2^Cre^* and *Six2^cre^Tet2^f/f^* mice subjected to sham-operation or UUO surgery. **(G)** Representative images of Sirius Red-stained kidney sections from *Six2^Cre^* and *Six2^cre^Tet2^f/f^* mice subjected to sham-operation or UUO surgery. (Right panel) Quantification (as a percentage of positive area) of Sirius Red-stained kidney sections from *Six2^Cre^* and *Six2^cre^Tet2^f/f^* mice subjected to sham-operation or UUO surgery. In the sham-operated group: *Six2^Cre^* (n = 4) and *Six2^cre^Tet2^f/f^* (n = 3). In the UUO-operated groups: n = 4 each. **(H)** Relative transcript level of *Fibronectin1* (*Fn1)*, *Collagen1a1* (*Col1a1)*, *Collagen1a1* (*Col3a1)*, *Interleukin 6* (*Il6*), *C-C motif chemokine ligand 2 (Ccl2)*, and *Vascular cell adhesion molecule 1 (Vcam1)* in the kidneys from *Six2^Cre^* and *Six2^cre^Tet2^f/f^* mice subjected to sham-operation or UUO surgery. In each of the sham-operated and UUO-operated groups: n = 6. **(I)** Representative Western blots and densitometric quantification of FN and alpha-smooth muscle (aSMA) in the kidneys of *Six2^Cre^* and *Six2^cre^Tet2^f/f^* mice subjected to sham-operation or UUO surgery. In each of the sham-operated and UUO-operated group: n = 3. **(J)** Experimental design for adenine-induced chronic kidney disease model. *Six2^Cre^* and *Six2^cre^Tet2^f/^*^f^ mice were placed on control or adenine diet. Mice were euthanized 4 weeks after starting the adenine food. **(K)** Serum BUN levels were measured in *Six2^Cre^* and *Six2^cre^Tet2^f/f^* mice on control or adenine diets. In the control diet-fed groups: n = 4 each. In adenine diet-fed groups: n = 5 each. **(L)** Representative images of H&E-stained kidney sections from *Six2^Cre^* and *Six2^cre^Tet2^f/f^* mice on control or adenine diet. **(M)** Representative images of Sirius Red-stained kidney sections from *Six2^Cre^* and *Six2^cre^Tet2^f/f^* mice on control or adenine diet. (Right panel) Quantification (as a percentage of positive area) of Sirius Red-stained kidney sections. In the control diet-fed groups: n = 3 each; In adenine diet-fed groups: n = 5 each. **(N)** Relative transcript level of *Fn1*, *Col1a1*, and *Col3a1* in the kidney samples from *Six2^Cre^* and *Six2^cre^Tet2^f/f^* mice on control or adenine diet. In the control diet-fed group: *Six2^Cre^* (n = 5) and *Six2^cre^Tet2^f/f^* (n = 4). In adenine diet-fed group: *Six2^Cre^* (n = 5) and *Six2^cre^Tet2^f/f^* (n = 6). **(O)** Representative Western blots and densitometric quantification of FN and aSMA in kidneys of *Six2^Cre^* and *Six2^cre^Tet2^f/f^* mice on control or adenine diet. In the control diet-fed groups: n = 3 in each. In adenine diet-fed groups: n = 4 each. \**p* < 0.05, \*\**p* < 0.01, \*\*\**p* < 0.001, \*\*\*\**p* < 0.0001 versus indicated group. Scale bar = 20 μm.

To understand the role of *TET2* in kidney fibrosis, we subjected both control and *Six2^Cre^Tet2^f/f^* mice to unilateral ureteral obstruction (UUO) injury (Figure 2E) and also analyzed their phenotypes in an adenine diet-induced chronic kidney fibrosis model (Figure 2J). Histological examination revealed more severe kidney fibrosis, including more dilated tubules with or without proteaceous cast, increased immune cell infiltration, and loss of brush border in kidneys of UUO surgery or adenine-diet-subjected *Six2^Cre^Tet2^f/f^* mice compared with those in UUO/adenine-subjected *Six2^Cre^* mice (Figure 2, F and L). Sirius Red staining indicated elevated profibrotic collagen deposition in kidneys of *Six2^Cre^Tet2^f/f^* mice with UUO surgery compared to kidneys of UUO/adenine-subjected control *Six2^Cre^* mice (Figure 2, G and M). Expression of fibrosis markers (*Fibronectin 1*, *Collagen1a1*, and *Collagen3a1*) was higher in kidneys of UUO/adenine-subjected *Six2^Cre^Tet2^f/f^* mice compared to *Six2^Cre^* mice with UUO surgery (Figure 2, H and N) and inflammation markers (*Interleukin 6*, *C-C motif chemokine ligand 2*, and *Vascular cell adhesion molecule 1*) was higher in kidneys of UUO-subjected *Six2^Cre^Tet2^f/f^* mice compared to kidneys of UUO-subjected *Six2^Cre^* mice (Figure 2H). Protein expression analysis by immunoblotting further confirmed that fibronectin1 and aSMA expressions were prominently increased in kidneys of UUO/adenine-subjected *Six2^Cre^Tet2^f/f^* mice compared to UUO-subjected *Six2^Cre^* mice (Figure 2, I and O).

As *Six2* is expressed during kidney development, we generated mice with conditional inducible genetic deletion of *Tet2* in tubule cells by crossing *Pax8^rtTA^* mice and *TRE^Cre^* mice with *Tet2^f/f^* mice (Figure 3A). Gene expression analysis confirmed the reduction in *Tet2* expression in the kidneys of *Pax8^rtTA^TRE^Cre^Tet2^f/f^* mice compared to*Tet2^f/f^* mice (Figure 3B). We observed that *Pax8^rtTA^TRE^Cre^Tet2^f/f^* mice were born at the expected Mendelian ratio and appeared healthy at birth, and we observed no differences in life span at 40 weeks of age. Most importantly, *Pax8^rtTA^TRE^Cre^Tet2^f/f^* were also protected from cisplatin-induced kidney injury (Figure 3, D-F) .

**Figure 3.**
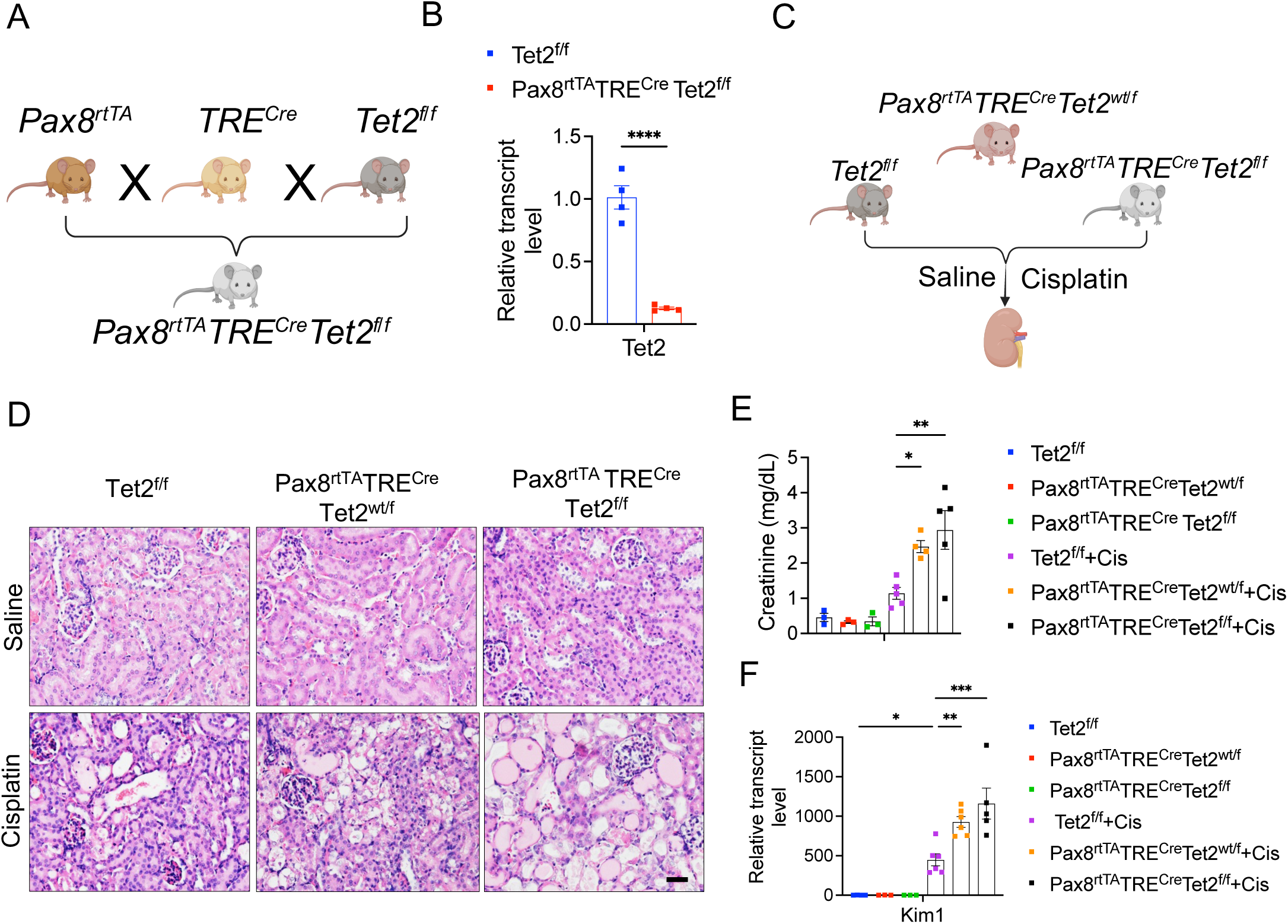
Tubule-specific *Tet2* deficiency exacerbates cisplatin-induced kidney injury. **(A)** Experimental scheme for the generation of *Pax8^rtTA^Cre^TRE^Tet2^f/f^* mice. **(B)** Relative transcript levels of *Tet2* in kidneys of control and *Pax8^rtTA^Cre^TRE^Tet2^f/f^* mice. N = 4 in each group. **(C)** Experimental design. *Tet2^f/f^*, *Pax8^rtTA^Cre^TRE^Tet2^wt/f^*, and *Pax8^rtTA^Cre^TRE^Tet2^f/f^* mice were injected with saline or cisplatin for 3 days. **(D)** Representative images of H&E-stained kidney sections from *Tet2^f/f^*, *Pax8^rtTA^Cre^TRE^Tet2^wt/f^*, and *Pax8^rtTA^Cre^TRE^Tet2^f/f^* mice, injected with saline or cisplatin. Scale bar = 20 μm. **(E)** Serum creatinine levels in *Tet2^f/f^*, *Pax8^rtTA^Cre^TRE^Tet2^wt/f^*, and *Pax8^rtTA^Cre^TRE^Tet2^f/f^* mice injected with saline or cisplatin. In the saline-injected groups: n = 3 each for each genotype. In the cisplatin-injected groups: *Tet2^f/f^*: n = 5, *Pax8^rtTA^Cre^TRE^Tet2^wt/f^*: n = 4, *Pax8^rtTA^Cre^TRE^Tet2^f/f^*: n = 5. **(F)** Relative transcript levels of *Kim1* in the kidneys of *Tet2^f/f^*, *Pax8^rtTA^Cre^TRE^Tet2^wt/f^*, and *Pax8^rtTA^Cre^TRE^Tet2^f/f^* mice injected with saline or cisplatin. In the saline-injected groups: n = 3 each for each genotype; In the cisplatin-injected groups: *Tet2^f/f^*: n = 6, *Pax8^rtTA^Cre^TRE^Tet2^wt/f^*: n = 6, *Pax8^rtTA^Cre^TRE^Tet2^f/f^*: n = 5. \**p* < 0.05, \*\**p* < 0.01, \*\*\**p* < 0.001 versus indicated group.

In summary, our data indicate that tubule-specific *Tet2* loss exacerbates kidney dysfunction both in AKI and CKD.

### Tet2 deficiency in tubule cells caused chromosome mis-segregation and accumulation of micronuclei

To investigate the mechanisms by which *Tet2* loss in tubule cells contributes to more severe kidney injury, we performed droplet-based snRNA-seq on kidney samples from wild-type (WT) and *Six2^Cre^Tet2^f/f^* mice at baseline and following UUO injury (Figure 4A). Following sequencing and alignment, we identified 38,714 high-quality cells. Subsequently, we applied dimensionality reduction using uniform manifold approximation and projection (UMAP) and graph-based clustering (Figure 4, B and C, and Supplemental Figure 5A). This analysis recognized 19 distinct clusters (Figure 4, B and C, and Supplemental Figure 5A), which were annotated based on the expression of previously published gene markers (Figure 4B) ^31^. *Tet2* level was markedly lower in *Six2^Cre^Tet2^f/f^* mice (Supplemental Figure 5C). We found that *Tet2* is mainly expressed in the proximal straight tubule (PST) (Supplemental Figure 5B) which was consistent with the in situ hybridization (ISH) and immunofluorescence (IF) staining data.

**Figure 4.**
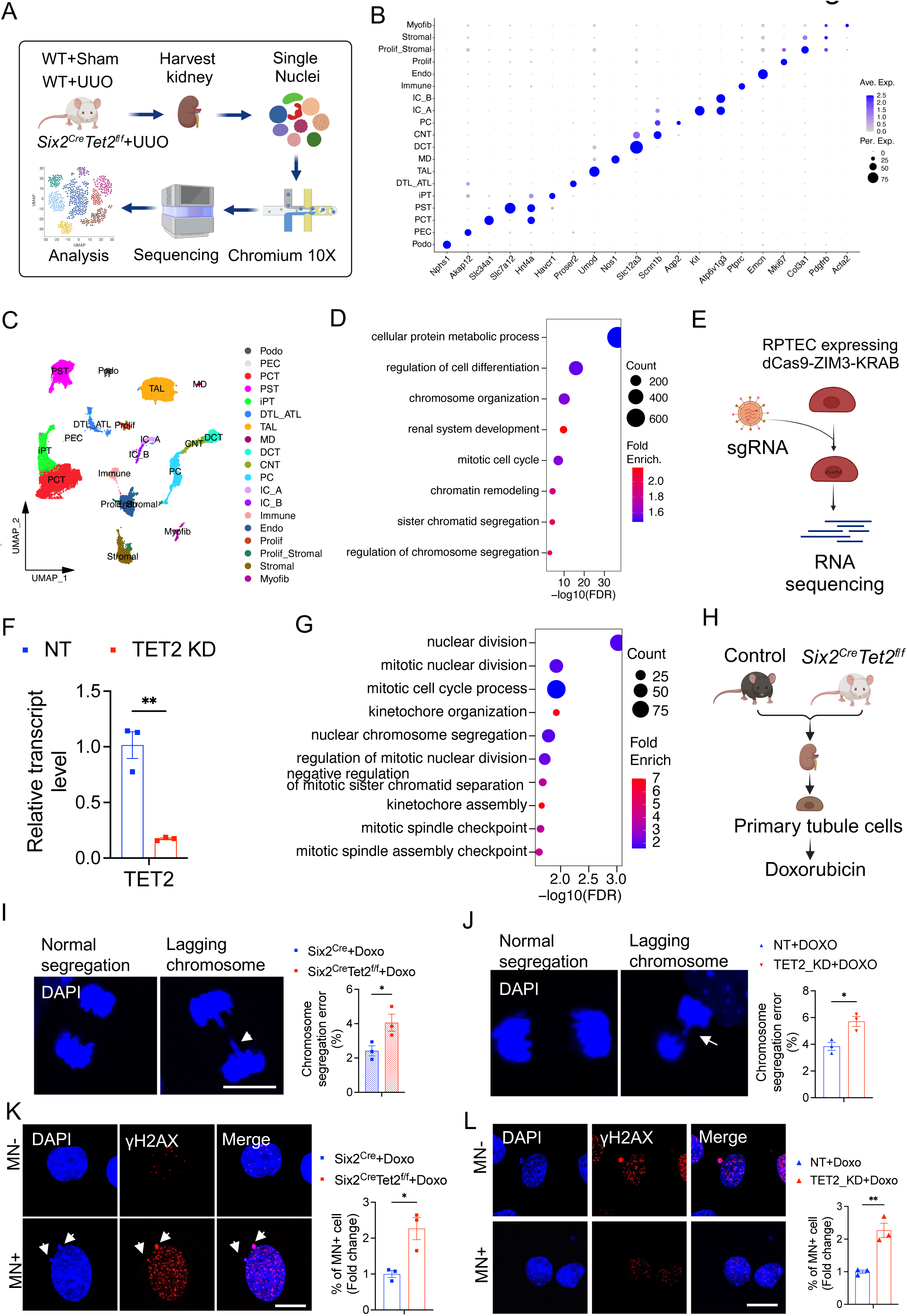
*Tet2* loss was associated with chromosome segregation defect and an increase in cytosolic micronuclei. **(A)** This diagram summarizes the process of nuclear isolation and single-nucleus RNA sequencing (snRNA-seq) of kidneys from wild type (WT) control and *Six2^Cre^Tet2^f/f^* mice subjected to sham-operation or UUO surgery, using 10X Genomics platform. **(B)** Bubble plots display the representative marker gene expression levels across 19 main cell clusters. Podocyte (Podo); Parietal epithelial cells (PEC); Proximal convoluted tubule (PCT); Proximal straight tubule (PST); Injured proximal tubule (iPT); Descending and ascending thin limb of loop of Henle (DTL_ATL); Thick ascending limb (TAL); Macular densa (MD); Distal convoluted tubule (DCT); Connecting tubule (CNT); Principal cells (PC); Type A intercalated cells of collecting duct (IC_A); Type B intercalated cells of collecting duct (IC_B); Endothelial cells (Endo); Proliferating cells (Prolif); Proliferating stromal (Prolif_Stromal); Myofibroblast (Myofib). The size of each bubble indicates the percentage of cells positiv for the marker, while the color intensity reflects the relative expression level. **(C)** UMAP dimension reduction of snRNA-seq of kidney samples from WT control and *Six2^Cre^Tet2^f/f^* mice subjected to sham-operation or UUO surgery. **(D)** GO functional analysis of differentially expressed genes (DEGs) observed in control WT and *Six2^Cre^Tet2^f/f^* mice performed with UUO surgery. Significantly enriched GO terms in biological process (*p* < 0.05) are presented, with *p*-values negative 10-base log transformed. The size indicates the number of differentially expressed genes in the group and color intensity indicates the fold enrichment change. DEGs, differentially expressed genes; GO, gene ontology. **(E)** This diagram summarizes the process of RNA sequencing (RNA-seq) process for control and *TET2* knockdown RPTEC. **(F)** Relative transcript levels of *TET2* in control and *TET2* knockdown RPTEC. n = 3 in each group. **(G)** GO functional analysis of genes with > 30% lower expression in *TET2* knockdown RPTEC compared to control cells. Significantly (*p* < 0.05) enriched GO terms in biological process are presented (p was negative 10-base log transformed). **(H)** Experimental design for the primary tubule cells isolated from kidneys of control and *Six2^Cre^Tet2^f/f^* mice and treated with doxorubicin. **(I-J)** Representative images of DAPI staining, showing chromosome segregation in control and *Tet2*-knockout primary tubule cells, as well as incontrol and *TET2-* knockdown RPTEC. (Right panel) Quantifications of the doxorubicin treated cells with chromosome segregation defect. n = 3 in each group. **(K-L)** Representative images of γH2AX and DAPI staining, showing primary tubule cells or RPTEC with or without micronuclei. (Right panel) Quantifications of the doxorubicin treated cells with micronuclei. In each group: n = 3. *p < 0.05, **p < 0.01 versus indicated group. Scale bar = 20 μm.

We next identified 2,218 differentially expressed genes (DEGs) in PST cells in kidneys from UUO-subjected *Six2^Cre^Tet2^f/f^* mice compared to UUO-subjected control mice kidneys (with an adjusted *p-*value < 0.05). Pathway analysis revealed significant enrichment in pathways related to metabolic processes, cell differentiation, and chromosome organization and segregation processes (Figure 4D).

To complement the mouse kidney analysis, we performed RNA sequencing (RNA-seq) analysis on control and *TET2* knockdown (KD) human renal proximal tubule epithelial cells (RPTECs) (Figure 4E). We generated these cells by transducing RPTECs, which stably express dCas9-ZIM3-KRAB, with either non-targeting control sgRNA or sgRNA targeting *TET2* (Figure 4E). We identified over 2,000 genes, including *TET2* (Figure 4F, Supplemental Figure 6), with a reduction of more than 30% in *TET2* KD RPTECs compared to control cells. Pathway analysis of these differentially expressed genes showed enrichment in processes related to nuclear division, mitotic cell cycle, and chromosome segregation (Figure 4G). We identified 45 genes involved in the chromosome segregation process, which were dramatically lowered in *TET2* KD cells, including *INO80* and *EME1* (Supplemental Table 1).

To confirm the role of *Tet2* in chromosome segregation in tubule cells, we next conducted in vitro studies by isolating primary tubule cells from the kidneys of *Six2^Cre^* control mice and *Six2^Cre^Tet2^f/f^* mice (Figure 4H), both in the presence and absence of doxorubicin, a DNA-intercalating agent (Figure 4H) ^32^. We observed various types of chromosome segregation errors in doxorubicin treated primary tubule cells isolated from kidneys of *Six2^Cre^Tet2^f/f^* mice compared to control cells (Figure 4I), including micronuclei formation^33,34^. There were more micronuclei in doxorubicin-treated tubule cells isolated from kidneys of *Six2^Cre^Tet2^f/f^* mice compared to those from kidneys of *Six2^Cre^* mice (Figure 4K). Similar results were obtained from control and *TET2* knockdown RPTECs treated with doxorubicin (Figure 4, J and L).

In summary, our unbiased gene expression and experimental analyses underscored the critical role of TET2 in chromosome segregation and prevention of micronuclei accumulation within kidney tubule cells.

### Loss of Tet2 in proximal tubule cell was associated with higher DNA damage due to impairment in DNA damage repair

Next, we analyzed RNA-seq data from a substantial collection (n>400) of both control and diseased human kidney samples. We focused on genes whose levels strongly correlated with *TET2*. We identified 160 genes, each with a false discovery rate (*FDR*) < 0.05, that correlated with *TET2* expression. Gene ontology analysis indicated strong enrichment for genes playing a role in DNA double-strand break (DSB) repair via homologous recombination (HR) (Supplemental Table 2). DNA DSBs are highly cytotoxic and must be repaired to maintain chromosomal integrity. To investigate the effects of *Tet2* loss on DNA damage in tubule cells, we measured DSB marker γH2AX level in cells treated with or without doxorubicin. We observed that γH2AX level was higher in doxorubicin-treated cells isolated from kidneys of *Six2^Cre^Tet2^f/f^* mice when compared to doxorubicin-treated cells isolated from kidneys of control mice (Figure 5A). Protein level of γH2AX was also elevated in kidneys of cisplatin-injected *Six2^Cre^Tet2^f/f^* mice or UUO surgery when compared to control cisplatin-injected mice or mice with UUO surgery (Figure 5, B and C).

**Figure 5.**
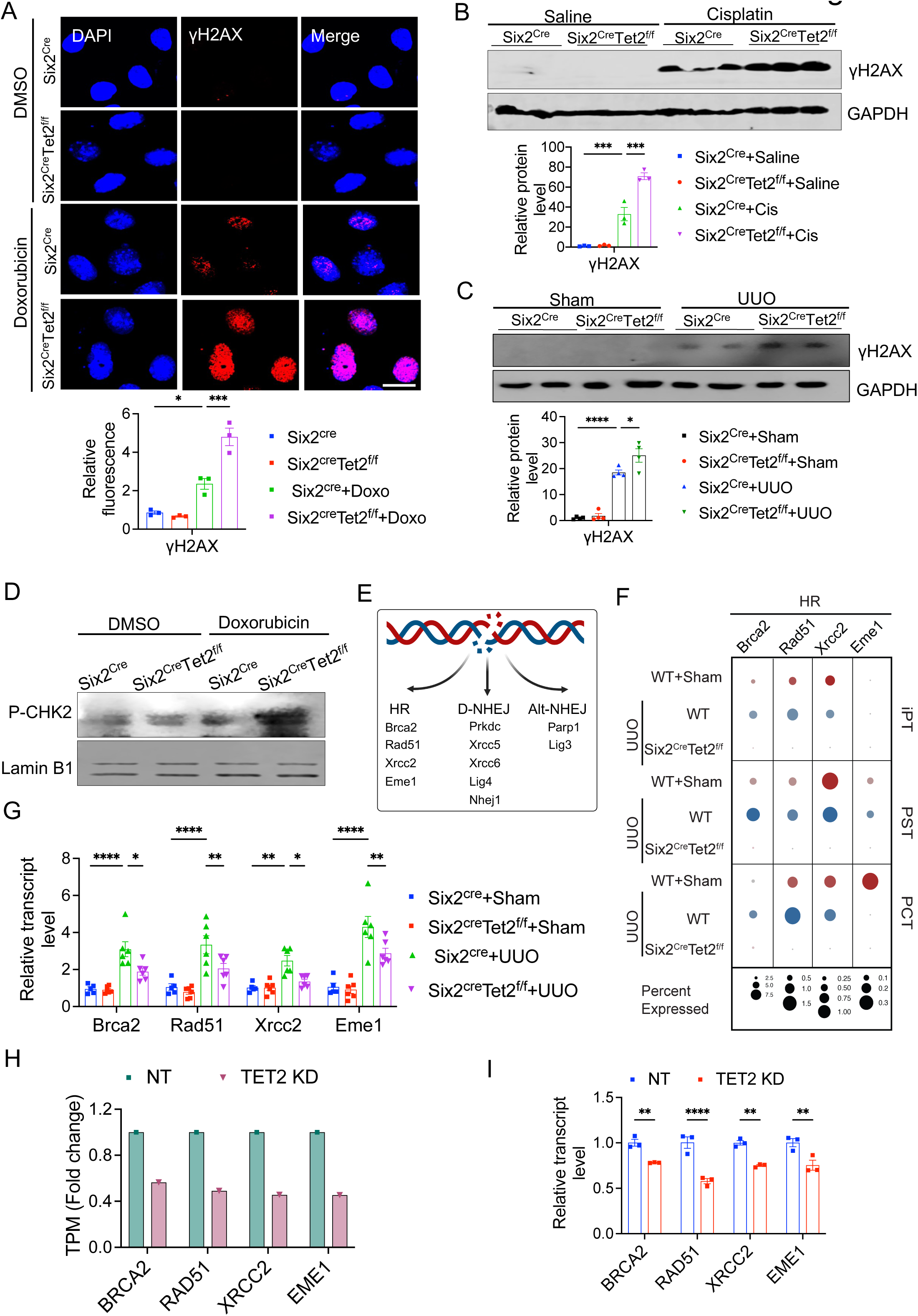
*Tet2* deficiency was associated with impaired DNA repair and accumulation of damaged DNA. **(A)** Representative images of immunofluorescence staining for γH2AX and DAPI in tubule cells isolated from control and *Six2^Cre^Tet2^f/f^* mice and treated with doxorubicin. n = 3 in each group. **(B-C)** Representative Western blots for γH2AX of kidney samples from control *Six2^Cre^* and *Six2^Cre^Tet2^f/f^* mice injected with saline or cisplatin and from *Six2^Cre^* and *Six2^Cre^Tet2^f/f^* mice subjected to sham-operation or UUO surgery. (Bottom panel) Densitometric quantification of γH2AX levels normalized to GAPDH. In the saline-injected or cisplatin-injected groups: n = 3 each. In the sham-operated or UUO-operated groups: n = 4 each. **(D)** Representative Western blots for nuclear p-CHK2 in tubule cells isolated from *Six2^Cre^* and *Six2^Cre^Tet2^f/f^* mice kidney and treated with or without doxorubicin. Lamin B1 is used as the loading control. **(E)** Scheme of genes involved in double strand break (DSB) repair pathways. **(F)** Bubble plots depicting the expression levels of *Brca2, Rad51, Xrcc2,* and *Eme1* across the PT clusters. The size of the circle represents the percentage of positive cells in the cluster, and the color of the bubble indicates the level of gene expression. **(G)** Relative transcript level of *BRCA2 DNA repair associated* (*Brca2)*, *RAD51 recombinase* (*Rad51), X-ray repair cross complementing 2* (*Xrcc2),* and *Essential meiotic structure-specific endonuclease 1 (Eme1)* in the kidneys obtained from *Six2^Cre^* and *Six2^cre^Tet2^f/f^* mice subjected to sham-operation or UUO surgery. *Gapdh* was used for normalization. In the sham-operated groups: *Six2^Cre^* (n = 5) and *Six2^cre^Tet2^f/f^* (n = 6); In the UUO-operated groups: n = 6 each. **(H)** Relative expression of *BRCA2*, *RAD51*, *XRCC2*, and *EME1* expression in control and *TET2*-knockdown RPTECs as determined by analysis of the RNA-seq dataset. **(I)** Relative transcript levels of *BRCA2*, *RAD51, XRCC2*, and *EME1* in control and *TET2*-knockdown RPTECs measured by qRT-PCR. n = 3 in each group. \**p* < 0.05, \*\**p* < 0.01, \*\*\**p* < 0.001, \*\*\*\**p* < 0.0001 versus indicated group. Scale bar = 20 μm.

Altered DNA damage response (DDR) during mitosis can lead to the accumulation of damaged DNA and chromosome mis-segregation ^32, 35^. We observed a higher level of phosphorylated checkpoint kinase 2 (p-CHK2), which is a marker of the DDR, in doxorubicin-treated cells isolated from the kidneys of *Six2^Cre^Tet2^f/f^* mice when compared with those in doxorubicin-treated control cells isolated the kidneys of *Six2^Cre^* mice (Figure 5D). These results are consistent with the observation of increased DNA damage and pronounced chromosome segregation errors in tubule cells lacking *Tet2*.

We hypothesized that damaged DNA repair might be impaired in absence of TET2, which would lead to damaged DNA accumulation. DNA DSBs are repaired by HR and non-homologous end repair (dominant D-NHEJ or alternative Alt-NHEJ) (Figure 5E) ^36^. We observed that the expression of HR pathway genes, including *BRCA2 DNA repair associated* (*Brca2*), *RAD51 recombinase* (*Rad51*), *X-ray repair cross complementing 2* (*Xrcc2*), and *Essential meiotic structure-specific endonuclease 1* (*Eme1*), was dramatically lower in PT cells of *Six2^Cre^Tet2^f/f^* mice compared to those in control PT cells from mice with UUO surgery (Figure 5F). Similar changes were observed in bulk gene expression data from the whole kidneys of *Six2^Cre^Tet2^f/f^* mice with UUO surgery (Figure 5G). In line with the snRNA-seq results, *TET2* KD RPTEC showed lower expression of HR pathway genes (Figure 5, H and I). D-NHEJ and Alt-NHEJ pathway genes, including *DNA activated, catalytic polypeptide (Prkdc), X-ray repair cross complementing 5* (*Xrcc5*), *X-ray repair cross complementing 1* (*Xrcc6*), *DNA ligase* 4 (*Lig4*), *Non-homologous end joining factor 1* (*Nhej1*), *Poly (ADP-ribose) polymerase 1* (*Parp1*), and *DNA ligase 3* (*Lig3*), were lower in PT cells of kidneys of *Six2^Cre^Tet2^f/f^* mice compared to kidneys obtained from control mice (Supplemental Figure 8A). However, we did not observe change in the expression of these genes in kidneys of *Six2^Cre^Tet2^f/f^* mice following UUO injury (Supplemental Figure 8, B and C) or in the *TET2* KD cells (Supplemental Figure 8, D-G).

Amongst the genes that showed consistent changes in PT cells of *Six2^Cre^Tet2^f/f^* mice and RPTECs with *TET2* loss was the *INO80 complex ATPase subunit* (*INO80)*. *INO80* has been proposed to play a role in DNA damage repair (both by HR and NHEJ pathways)^37–41^. Most importantly, *INO80* is located in a genomic region association with eGFR, as indicated by eGFR GWAS (rs4924532, *P* = 2.96 × 10^-40^) (Figure 6A). Furthermore, GWAS annotation with kidney tubule eQTL (rs4924532, *P* = 6.45×10^-6^) data indicated that *INO80* is the target of the eGFR GWAS variants in kidney tubule cells (Figure 5A). *Ino80* expression was elevated in the PT cells of UUO injured mouse kidneys (Figure 6D).

**Figure 6.**
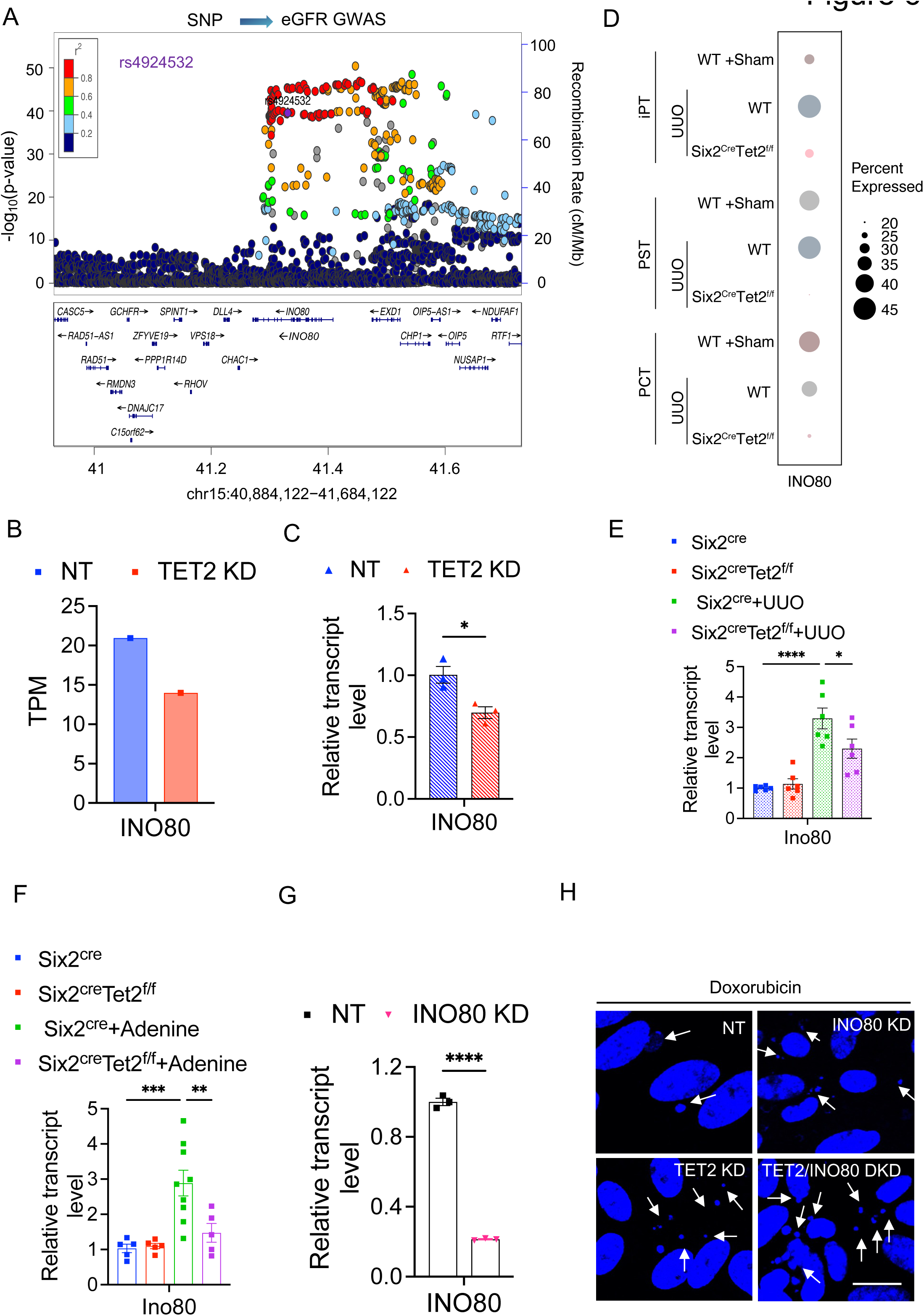
*Ino80* is another kidney disease risk gene associated with impaired DNA damage repair. **(A)** LocusZoom plot of GWAS (genotype and eGFRcrea association, n = 1,508,659), **(B)** RNA-seq of control and *TET2*-knockdown RPTECs showing *INO80* expression in these cells. **(C-F)** Relative transcript levels of *INO80* in control and *TET2*-knockdown RPTECs, and its levels in kidney samples from *Six2^Cre^* and *Six2^Cre^Tet2^f/f^* mice subjected to sham-operation or UUO surgery or on control or adenine diets. For the control and *TET2*-knockdown (KD) RPTEC groups, n = 3 for each groups. In the sham-operated and UUO-operated groups: n = 6 for each group. In the control diet-fed groups: n = 5 each. For the adenine diet-fed group, *Six2^Cre^* mice had n = 9, and *Six2^cre^Tet2^f/f^* mice had n = 5. **(G)** Relative transcript levels of *INO80* in *INO80*-knockdown cells compared to control cells, with n= 3 in each group. **(H)** Representative images of immunofluorescence staining showing micronuclei (MN) in control, *INO80-*knockdown, *TET2-* knockdown, and *TET2* and *INO80* double knockdown RPTECs. *p < 0.05, **p < 0.01, ***p < 0.001, ****p < 0.0001 versus indicated group. Scale bar = 20μm.

Similarly, an increased number of micronuclei were observed in *INO80* knockdown RTPEC following doxorubicin treatment, indicating the role of *INO80* in DDR (Figure 6H).

In summary, these results indicate that *Tet2* loss leads to impaired DNA repair and accumulation of DNA damage and chromosome segregation defects in kidney tubule cells.

### Role of cGAS-STING activation in Tet2 loss-induced kidney disease development

We next hypothesized that activation of cytosolic nucleotide sensors (cGAS/STING) might mediate the deleterious effect of *TET2* loss, as cytosolic DNA and micronuclei are potent activators of this pathway ^33^. The *TET2* KD RPTEC showed higher cGAS-STING levels (Figure 7A). Expression of downstream STING target genes, including *Radical S-adenosyl methionine domain containing 2* (*RSAD2*), *2’-5’-oligoadenylate synthetase 1* (*OAS1*), and *MX dynamin like GTPase 1* (*MX1*), were also higher in *TET2* knockdown cells (Supplemental Figure 10A), which we confirmed by qRT-PCR (*Isg15*, *Isg20*, *Ifit1*, *Ifit2*, *Ifit3*, and *Mx1*) (Figure 7B). Protein level of cGAS, STING, phosphorylated TBK1 (p-TBK1), phosphorylated IRF3 (p-IRF3), and phosphorylated P65 (p-P65) was higher in cisplatin-injected *Six2^Cre^Tet2^f^*^/f^ mice compared to cisplatin-treated control *Six2^Cre^* mice (Figure 7C and Supplemental Figure 10B). Immunofluorescence staining showed that p-STING level was higher in kidneys of *Six2^Cre^Tet2^f/f^* mice injected with cisplatin compared to those of *Six2^Cre^* mice (Supplemental Figure 10C). ISH staining showed higher *Isg15* expression in kidneys of *Six2^Cre^Tet2^f/^*^f^ mice injected with cisplatin when compared to those of control *Six2^Cre^* mice injected with cisplatin (Figure 7D). Similar results were obtained in the UUO disease model (Figure 7D and Supplemental Figure 10D).

**Figure 7.**
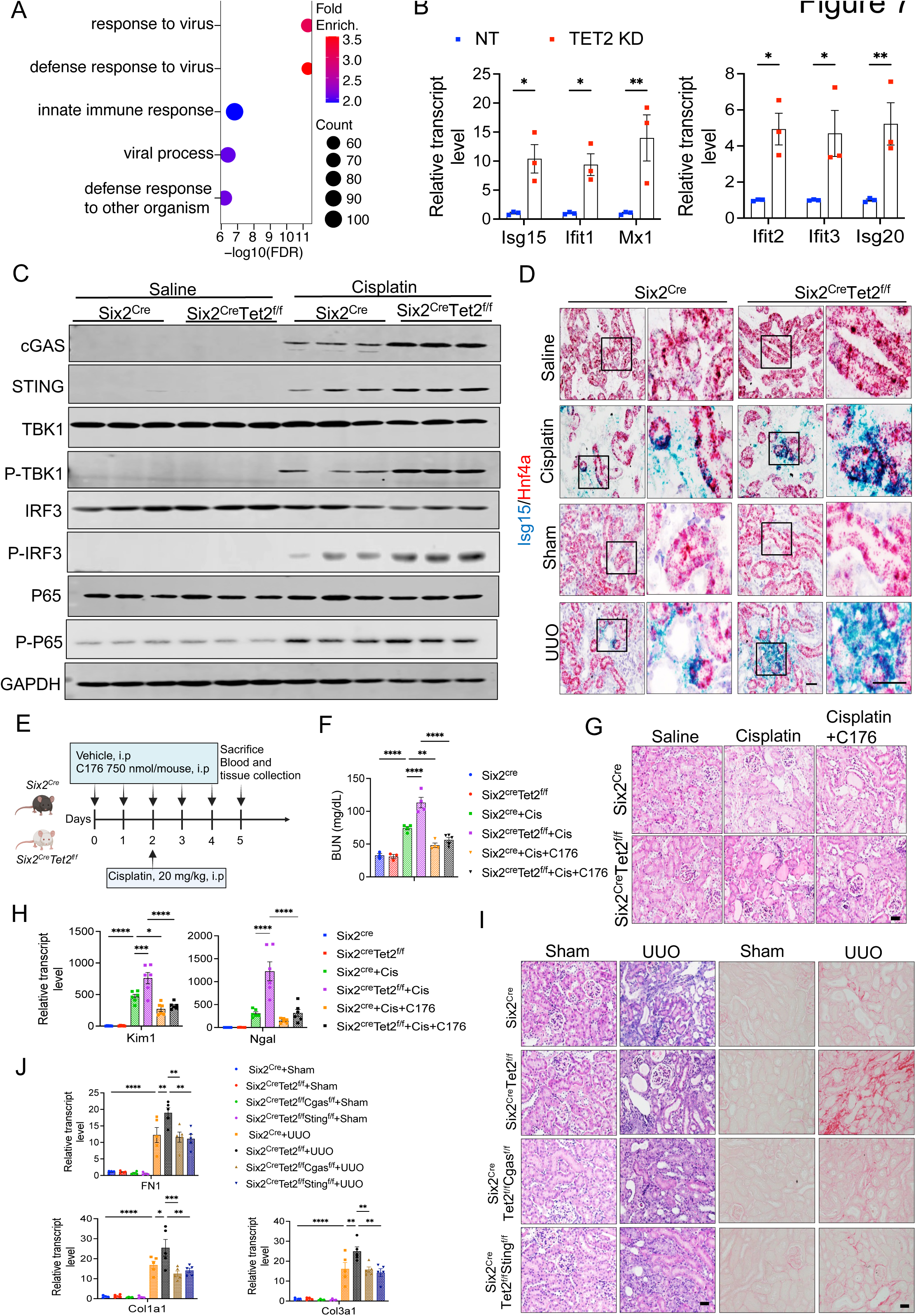
cGAS and STING mediate the *TET2 -*loss induced kidney damage. **(A)** GO functional analysis of genes with levels more than 1.5-fold in *TET2*-knockdown RPTECs compared to control cells. Significantly enriched GO terms in biological process (*p* < 0.05) are presented, with p-values presented as negative base-10 log-transformed. **(B)** Relative transcript levels of *Isg15, Ifit1, Ifit2, Ifit3, Mx1*, and *Isg20* in control and *TET2* knockdown RPTECs. In each group: n = 3. **(C)** Representative Western blots for *cGAS, STING, TBK1, p-TBK1, IRF3, p-IRF3, P65,* and phosphorylated p65 (p-p65) in kidney samples from *Six2^Cre^* and *Six2^Cre^Tet2^f/f^* mice subjected to sham operation or UUO surgery. **(D)** Representative in situ hybridization images of *Isg15* (blue) and *Hnf4a* (red) in the kidney sections from *Six2^Cre^* and *Six2^Cre^Tet2^f/f^* mice, either injected with saline or cisplatin or subjected to sham operation or UUO surgery. **(E)** Experimental design. *Six2^Cre^* and *Six2^Cre^Tet2^f/f^* mice injected with saline or cisplatin in the presence or absence of the STING inhibitor C176. **(F)** Serum BUN levels in *Six2^Cre^* and *Six2^Cre^Tet2^f/f^* mice injected with saline or cisplatin either in the presence or absence of C176.For the saline-injected groups, n = 3 for each group. For the cisplatin-injected groups, n = 4 for each group. In the cisplatin and C176-injected groups: *Six2^Cre^* (n = 4) and *Six2^cre^Tet2^f/f^* (n = 5). **(G)** Representative images of H&E-stained kidney sections from *Six2^Cre^* and *Six2^Cre^Tet2^f/f^* mice injected with saline or cisplatin either in the presence or absence of C176. **(H)** Relative transcript level of *Kim1* and *Ngal* in kidney samples from *Six2^Cre^* and *Six2^Cre^Tet2^f/f^* mice injected with saline or cisplatin, either in the presence or absence of C176. In the saline-injected groups: n = 4 each. In the cisplatin-injected groups: n = 5 each. In the cisplatin and C176-treated groups: n = 5 each. **(I)** Representative images of H&E-stained and Sirius Red-stained kidney sections from *Six2^Cre^*, *Six2^Cre^Tet2^f/f^*, *Six2^Cre^Tet2^f/f^Cgas^f/f^*, and *Six2^Cre^Tet2^f/f^Sting^f/f^* mice subjected to sham-operation or UUO surgery. **(J)** Relative transcript levels of *Fn1, Col1a1,* and *Col3a1* in kidney samples from *Six2^Cre^*, *Six2^Cre^Tet2^f/f^*, *Six2^Cre^Tet2^f/f^Cgas^f/f^*, and *Six2^Cre^Tet2^f/f^Sting^f/f^* mice subjected to sham-operation or UUO surgery. In the sham-operated groups: n = 4 each; In the UUO-operated groups: n = 5 each. .\**p* < 0.05, \*\**p* < 0.01, \*\*\**p* < 0.001, \*\*\*\**p* < 0.0001 versus indicated group. Scale bar = 20 μm.

To prove that cGAS-STING activation mediates kidney injury induced by tubule-specific loss of *Tet2*, we examined the effect of the STING inhibitor C176 in cisplatin-induced AKI mice (Figure 7E). The expression of STING target gene, such as *Isg15,* was lower in kidneys of *Six2^Cre^Tet2^f/f^* mice treated with C176, indicating the effectiveness of the drug (Supplemental Figure 11). BUN levels, a marker kidney function, were lower in C176-treated mice compared to those in cisplatin-treated *Six2^Cre^Tet2^f/f^* mice (Figure 7F). Histological examinations showed less severe kidney injury (such as dilated tubules and loss of brush border) in kidneys of *Six2^Cre^Tet2^f/f^* mice treated with both cisplatin and C176, compared to those treated with cisplatin alone (Figure 7G). Levels of kidney injury markers, *Kim1* and *Ngal*, were lower in C176-injected animals (Figure 7H).

Finally, we used a genetic approach to prove the role of cGAS and STING in TET2-mediated kidney injury. We crossed *Six2^Cre^* mice and *Tet2^f/f^* mice with *Cgas^f/f^* mice or *Sting^f/f^* mice (Supplemental Figure 12A) and then subjected these animals to UUO injury (Supplemental Figure 12B). Protein levels of cGAS and STING in the kidneys confirmed the knockout efficiency of cGAS in *Six2^Cre^Tet2^f/f^Cgas^f/f^* mice and STING in *Six2^Cre^Tet2^f/f^Sting^f/f^* mice when compared to kidneys of UUO subjected *Six2^Cre^Tet2^f/f^* mice (Supplemental Figure 12C). Histological analyses revealed reduced injury levels in kidneys of *Six2^Cre^Tet2^f/f^Cgas^f/f^* and *Six2^Cre^Tet2^f/f^Sting^f/f^* mice subjected to UUO surgery, compared to *Six2^Cre^Tet2^f/f^* mice also undergoing UUO surgery (Figure 7I). Sirius Red staining indicated less collagen accumulation in kidneys of UUO-subjected *Six2^Cre^Tet2^f/f^Cgas^f/f^* mice and *Six2^Cre^Tet2^f/f^Sting^f/f^* mice compared to those in *Six2^Cre^Tet2^f/f^* mice with UUO surgery (Figure 7I and Supplemental Figure 12D). Transcript levels of fibrosis markers, including *Fibronectin1*, *Collagen1a1*, and *Collagen 3a1,* were lower in kidneys of UUO-subjected *Six2^Cre^Tet2^f/f^Cgas^f/f^* and *Six2^Cre^Tet2^f/f^Sting^f/f^* mice compared to those in *Six2^Cre^Tet2^f/f^* mice subjected to UUO (Figure 7J). The protein level of fibronectin 1 was lower in the kidneys of UUO-subjected *Six2^Cre^Tet2^f/f^Cgas^f/f^* and *Six2^Cre^Tet2^f/f^Sting^f/f^* mice compared to those in *Six2^Cre^Tet2^f/f^* mice subjected to UUO (Supplemental Figure 12C).

In summary, these findings underscore the critical role of the cGAS-STING in mediating *Tet2* loss-induced kidney injury and kidney disease development.

## DISCUSSION

Here, we demonstrate that *TET2* is a kidney disease risk gene using a multi-omics integration of kidney function GWAS fine mapping, human kidney meQTL, eQTL data, single-cell chromatin accessibility annotation, and CRISPR/Cas9-mediated genome editing. Mice and cells with genetic deletion of *TET2* in kidney tubule cells showed impaired DNA damage repair, resulting in the accumulation of cytosolic micronuclei, downstream cGAS-STING activation, and the development of kidney fibroinflammation.

Loss of function somatic mutations in TET2 within blood cell progenitors have recently gained major scientific interest as they are commonly observed in older individuals. Clonal hematopoiesis of indeterminate potential (CHIP) is a common aging-related phenomenon in which early blood cell progenitors contribute to the formation of a genetically distinct subpopulation of blood cells. Somatic mutations most commonly occur in *TET2* and *DNMT1*. CHIP was first reported to be associated with myeloid malignancies^42^, but later studies have uncovered the association of CHIP with cardiovascular disease ^43^, melanoma progression^44^, and Parkinson‘s disease ^45^. The role of CHIP and somatic *TET2* mutations in kidney disease development has also been proposed ^46, 47^. Our studies suggest that germline regulatory genetic variants in TET2 predispose individuals to kidney disease development. Future studies shall determine the role of CHIP and *TET2* loss in blood cell progenitors, as well as the potential interaction between tubule cells and hemopoietic stem cells in kidney disease.

TET2 is broadly expressed in the body. It is interesting to note that *TET2* is almost exclusively expressed in the S3 segment of the PT in both mouse and human kidneys. Our study indicates that germline genetic variants at the *TET2* locus will alter tubule-specific *TET2* levels. Mice with a global Tet2 deletion exhibited exacerbated acute renal injury following cisplatin injection ^48^ and ischemia-reperfusion injury ^49^. The role of TET2 in diabetic kidney disease has been proposed, based on findings from cell culture studies^50^. Our study is the first to utilize mice with a kidney/tubule-specific deletion of *Tet2,* specifically exploring its role in the tubule cells, furthermore we show that the effect is not related to developmental programming as mice with conditional inducible deletion of

TET2 in mature tubule cells showed the same phenotype, providing direct evidence of how germline genetic variations can alter*Tet2* levels in the kidney tubule.

Our studies indicate the important role of impaired DNA damage repair, likely via the HR pathway in kidney disease development. Future studies shall examine exact molecular pathways regulated by TET2. Our findings are consistent with a prior study showing that *Tet2* regulates HR in leukemia cells ^51^. TET2 has also been localized to DSB in HeLa cells, and human embryonic stem cells and critical for maintain genomic integrity ^52^. DNA methylation, particularly at CpG islands within promoter regions, can regulate the expression of key genes involved in the DDR. Alterations in methylation patterns can either impair or enhance DDR.

The role of DDR in kidney disease goes beyond *TET2*, as we also identified *INO80* as another gene related to DNA repair and associated with kidney disease risk. The INO80 complex is a chromatin remodeling complex that regulates gene expression, DNA repair, and replication. Our integrated GWAS and follow-up prioritization tool identified 48 kidney disease risk genes playing a role in DDR, including *BRCA2, NIMA related kinase 4* (*NEK4*)*, and O-6-methylguanine-DNA methyltransferase* (*MGMT*) ^7^. Overall, our study suggests DDR plays a critical and broader role in kidney disease development. Aging is the strongest risk factor for kidney disease development however the exact mechanism of aging-associated kidney disease is poorly understood. It is known that DNA methylation degrades with aging, and increased damage DNA accumulation and altered DDR are observed during aging. Future studies shall establish the role of methylation changes and DDR in aging associated kidney disease development.

Our preliminary analysis indicates that TET2 effect is linked to its enzymatic function. First, we observed important differences in methylation cytosine levels between control, KO mice and those in a disease state. Our prior studies indicated that Tet2, in conjunction with Tet3, markedly alters kidney development by interfering with the methylation of WNT pathway genes. Finally, we found that vitamin C, which can enhance the enzymatic activity of TET enzymes, ameliorated kidney disease development in mice with lower TET2 levels. Future studies shall analyze both methylation and hydroxymethylation levels in the Tet2 knockout animals.

Our studies indicated that the *Tet2* loss results in accumulation of micronuclei in tubule cells, activating the cGAS-STING pathway. We found that damaged DNA accumulating in injured kidney tubule cells leads to alterations in chromosome segregation, including delays, which is likely the cause of micronuclei accumulation. Cytosolic DNA, including micronuclei, has been shown to activate the cytosolic DNA sensors, such as cGAS and STING, in both aging and cancer cells. To demonstrate the critical role of cGAS and STING in tubule cells, we generated mice with a tubule-specific genetic deletion of cGAS and STING. These mice showed protection from kidney disease, indicating the critical role of this pathway in DDR alterations and kidney disease development.

Finally, we show that pharmacological inhibition of STING protected mice from severe disease, indicating that the development of effective STING inhibitors might be beneficial for patients with kidney disease specifically in patients carrying DDR pathway gene variants.

In conclusion, here we identify *TET2* as a kidney disease risk gene, which plays an important role in DNA damage repair and preventing altered chromosome segregation, cytosolic micronuclei accumulation, and cGAS-STING activation in kidney tubule cells.

## METHODS

### Animal studies

Mouse studies were approved by the Institutional Animal Care and Use Committee at the University of Pennsylvania. The *Tet2^f/f^* (stock #017573), *Six2^Cre^* (stock # 009606), *Pax8^rtTA^* (stock #007176), TetO-Cre (*TRE^Cre^*) (stock #00623), and *Sting^f/f^* (stock #031670) mice were purchased from the Jackson Laboratory. *Cgas ^f/f^* mice were kindly provided by C.Rice (Rockefeller University)^53^. *Tet2^f/f^* was crossed with *Six2^Cre^* mice to generate *Six2^Cre^Tet2^f/f^* mice. *Pax8^rtTA^* and *TRE^Cre^* mice were crossed with *Tet2^f/f^* to generate *Pax8^rtTA^TRE^Cre^Tet2^f/f^* mice. *Cgas ^f/f^* and *Sting ^f/f^* were crossed with *Six2^Cre^Tet2^f/f^* mice to generate *Six2^Cre^Tet2^f/f^*Cgas *^f/f^* mice and *Six2^Cre^Tet2^ff/^* Sting *^f/f^* mice, respectively. For the cisplatin injury model, 6-8-week-old mice were intraperitoneally injected with cisplatin (20 mg/kg; dissolved in 0.9% saline) and euthanized on day 3. For the unilateral ureteral obstruction model, 6-8-week-old mice underwent ligation of the left ureter and were euthanized on day 4. For adenine mouse disease model, 8-10-week-old mice were given a control (0.6% calcium and 0.9% phosphorus adjusted diet) or an adenine (0.2% adenine, 0.6% calcium and 0.9% phosphorus) diet and euthanized 4 weeks after starting the adenine food. In the STING pharmacological inhibition experiment, C176 (750 nm per mouse, Bio Vision, #B2341) or DMSO dissolved in 85 μl corn oil was administered intraperitoneally. This administration occurred at 2 h, 24 h, and 48 h before the injection of cisplatin (20 mg/kg) or saline, and then daily after cisplatin injection. All mice were provided food and water ad libitum and monitored daily.

### Single nucleus RNA sequencing

Nuclei were extracted from flash-frozen kidney tissue. Briefly, 10-30 mg frozen kidney tissue were minced on ice into pieces in lysis buffer (Tris-HC, NaCl, MgCl2, NP40, and RNAse inhibitor), transferred into a gentleMACS C tube and homogenized using gentleMACS homogenizer. Next, the homogenized tissue was filtered through a 40 μm strainer (Fisher Scientific, #08-771-1) and then centrifuged at 500 × g for 5 mins at 4°C. The pellet was resuspended in wash buffer (BSA, RNAse inhibitor, 1× PBS) and then filtered through a 40 μm Flowmi cell strainer (Sigma Aldrich, #BAH136800040-50EA).

After checking for nuclear quality and counting, 30,000 cells were loaded into the 10X Chromium Controller (PN-120223). The cDNA libraries were constructed using the 10X Chromium library construction kit (PN-1000190). Libraries were subjected to quality control using the Agilent Bioanalyzer High Sensitivity DNA kit (Agilent Technologies, 5067-4626). The following demultiplexing was used to sequence libraries using the Illumina Novaseq 6000 system with 2 × 150 paired-end kits: 28 bp Read1 for cell barcode and UMI, 10 bp i7 index, 10 bp i5 index, and 90 bp Read2 for transcript.

### Cell culture and lentiviral production

Primary tubular epithelial cells were isolated from kidneys of 3-4-week-old mice. Briefly, kidneys were minced into pieces and digested in RPMI 1640 medium (Gibco, #21875-034) containing collagenase IV (2 mg/ml, Sigma Aldrich) for 30 min at 37°C. Next, fetal bovine serum (FBS) was used to deactivate collagenase IV activity and cells were sequentially sieved through 100, 70, and 40 μm mesh. After removing red blood cells, the tubules cells were cultured in RPMI 1640 supplemented with 10% FBS, rhEGF (20 ng/ml), 1× insulin-transferrin-selenium (ITS), and 1% penicillin-streptomycin.

Immortalized human RPTECs (ATCC; CRL-4031) were cultured in Dulbecco’s modified Eagle’s medium (DMEM)-F12 supplemented with hydrocortisone (25 ng/ml), ascorbic acid (3.5 µg/ml), sodium selenite (8.65 ng/ml), transferrin (5.0 µg/ml), insulin (5.0 µg/ml), triiodo-L-thyronine (5 pM), prostaglandin E1 (25 ng/ml), rhEGF (10 ng/ml), sodium bicarbonate (1.2 mg/ml), and 1% penicillin-streptomycin, and G418 (0.1 mg/ml).

For the cytosolic micronuclei analysis experiments, the primary epithelial tubule cells or RPTECs were exposed to doxorubicin (1 μM) for 25 mins. For the DNA damage analysis experiments, the primary epithelial tubule cells or RPTECs were exposed to doxorubicin (1 μM) for 25 mins and analyzed after 8 h. For the cell death analysis experiments, the primary primary epithelial tubule cells were treated with doxorubicin (1 μM) for 24 h.

Human embryonic kidney (HEK) 293T (HEK293T) cells (ATCC, #CRL-3216), used for packaging lentivirus. They were cultured in DMEM (Mediatech, #MT10-013-CV) supplemented with 10% FBS and 1% penicillin-streptomycin. All cultured cells were maintained in a humidified 5% CO_2_ atmosphere at 37 °C.

For lentivirus production, HEK293T cells were transfected with the plasmid of interest, along with the lentiviral packing and envelope plasmids, pPAX2 (Addgene, #12260, a gift from Didier Trono) and pMD2.G (Addgene, #12259, a gift from Didier Trono), using polyethylenimine (PEI, Biotechne, #7854).

### Cell line generation

CRISPRi RPTEC cells were generated by stably transducing RPTECs with lentivirus expressing ZIM3 KRAB-dCas9-mCherry under the UCOE-SFFV promoter. Cells stably expressing mCherry were sorted using fluorescence activated cell sorting. The plasmid pHR-UCOE-SFFV-dCas9-mCherry-ZIM3-KRAB was a gift from Mikko Taipale (Addgene plasmid #154473). CRISPRi RPTEC cell line was verified by monitoring mCherry fluorescence over several generations. Knockdown was confirmed using functional analysis.

### CRISPR/Cas9 mediated genomic region deletion

All annealed sgRNA oligos were subcloned into a Cas9 expression plasmid (lentiCRISPR v2, Addgene, #52961, a gift from Feng Zhang) with the Bsmb1 site. All constructs were verified by Sanger sequencing.

Lentivirus was prepared as previously described. Lentivirus-containing supernatants supplemented with polybrene (Santa Cruz Biotechnology, #sc-134220) were used for lentiviral transduction for the human RPTEC. After 72 hours, cells were harvested, and *TET2* and *PPA2* expression was determined by qRT-PCR. At the same time, DNA was isolated, and sgRNAs target region deletion were determined by Sanger sequencing. sgRNA sequences are listed in Supplemental Table 3. The PCR primer sequences are listed in Supplemental Table 4.

### CRISPR interference

The sgRNA expressing plasmids were generated as previously published ^54^. Briefly, annealed sgRNA oligos were subcloned into LRG2.1 (Addgene #108098, a gift from Christopher Vakoc) with the Bsmb1 site. All constructs were verified by Sanger sequencing. The sgRNA sequences used in this study are in Supplementary Table 3.

Lentivirus was prepared as previously described. The lentivirus-containing supernatants, supplemented with polybrene, were immediately used for lentiviral transduction of human RPTEC stable expressing dCas9-KRAB-ZIM3.

### Human RPTEC RNA Sequencing

RNA was extracted using the RNeasy Mini Kit (Qiagen, Cat# 74106). After quality assessment by Agilent Bioanalyzer 2100, RNA-Seq libraries were generated using the TruSeq RNA Sample Prep Kit (Illumina, San Diego, CA). Low-quality bases were trimmed using Trim-galore, and RNA-seq reads were aligned to the human genome (hg19) using STAR (v2.6.1a) based on GENCODE v29 annotations. RSEM (v1.3.1) was used to quantify gene-level read counts and expressed as transcripts per million (TPM).

### RNA extraction and qRT-PCR

RNA was isolated using Trizol reagent (Invitrogen) and was transcribed into cDNA using cDNA Reverse Transcription Kit (Applied Biosystems, Cat. #4368813). qRT-PCR was performed using the SYBR Green Master Mix (Applied Biosystems, Cat. #A25742-PEC). The data were normalized and analyzed using the 2-ΔΔCt method with indicated reference gene. Primer sequences are provided in Supplemental Table 5.

### Western blot

Kidney tissue or cultured cells were homogenized in lysis buffer and were separated by SDS-PAGE and then transferred to PVDF membranes. Transferred blots were blocked for 1h in 5% non-fat milk or 5% BSA in Tris-buffered saline. Membranes were incubated with specific primary antibodies at 4°C overnight, followed by incubation with a horseradish peroxidase-conjugated anti-mouse antibody (1:5,000), or anti-rabbit antibody (1:5,000) at room temperature for 1 h. Resulting immunoblots were visualized using ECL Western Blotting Substrate in a LI-COR chemiluminescence imager (LI-COR), according to the manufacturers’ instructions.

### Histological analysis

Kidneys were fixed in 10% neutral formalin. Paraffin-embedded sections were stained with hematoxylin and eosin (H&E). Sirius Red staining was performed to determine the degree of fibrosis. *In situ* hybridization was performed using paraffin-embedded kidney tissue samples and the RNAscope 2.5 HD Duplex Detection Kit (bio-techne, Cat. #322436) following manufacturers’ original protocol. The following probes were used for the RNAscope assay: Mm-*Tet2* (bio-techne, Cat. # 511591), Mm-I*sg15* (bio-techne, Cat# 559271-O1), Mm-*Hnf4a* (bio-techne, Cat# 497651-C2). Immunofluorescence analyses were performed on paraformaldehyde-fixed cells and formalin-fixed, paraffin-embedded kidney sections. To visualize the expression of proteins in the kidney, paraffin-embedded sections were deparaffinized, rehydrated, and incubated with indicated primary antibodies. Slides were incubated with fluorescent conjugated secondary antibodies, counterstained and mounted with DAPI for nuclear stain. To visualize the protein expression in the paraformaldehyde-fixed cells, we permeabilized the cells with 0.2% Triton X 100, blocked with 3% BSA, then incubated with indicated primary antibodies and fluorescent conjugated secondary antibodies, and counterstained with mounted with DAPI.

### BUN and creatinine level

Blood urea nitrogen level (BUN) was measured using Infinity^TM^ Urea Liquid Stable Reagent (Pointe Scientific, #B7552150). Serum creatinine was detected by Creatinine Enzymatic and Creatinine Standard (DIAZYME #DZ072B-KY1). Both measurements were performed following the manufacturers’ instructions.

### Cytotoxicity assays

Mouse primary tubular epithelial cells were plated in a 96-well plate and treated with or without cisplatin for indicated time. LDH release was quantified using CytoTox 96 Non-Radioactive Cytotoxicity Assay (Promega #G1780) according to the manufacturer’s instructions. Briefly, medium was collected and incubated in a 96-well plate with the CytoTox reagent. After adding the stop solution, the absorbance signal was detected at 490 nm in a plate reader.

### Bioinformatic analysis for single-nucleus RNA sequencing data

We generated a gene-by-cell count matrix by aligning its fastq files to the mm10 reference dataset using 10X Genomics Cell Ranger ARC v2.0.2). The quality control (QC) included removing ambient mRNA contamination, eliminating low-quality cells and genes, and excluding doublet-like cells. We used “SoupX” to remove ambient mRNA. We defined low-quality cells as those with the following criterias: unique molecular identifiers (UMI) ≤ 500 or ≥ 10,000, the number of detected genes ≤300 or ≥ 3000, the percentage of mitochondrially encoded genes reads > 1%, complex ≥ 0.25. We further removed genes that were expressed in < 10 cells. “DoubletFinder” was used to exclude doublet-like cells. To process the data, we used Seurat 4.0.4.

The data were normalized and scaled using the NormalizeData() and ScaleData() functions in Seurat, and variable features were identified with FindVariableFeatures(). Cluster identities were assigned using expression of known marker genes ^31^. Differentially expressed genes (DEGs) in each cluster were determined using the FindAllMarkers() function with the “*<u>MAST”</u>* test for transcripts present in at least 10% of cells with an absolute log2-fold-change (log2FC) threshold of 0.25. A Bonferroni adjusted *P*-value (Padj) < 0.05 was considered statistically significant.

### Quantification and statistical analysis

**S**tatistical analyses were performed using GraphPad Prism software (GraphPad Software Inc., La Jolla, CA). A two-tailed t-test was used to compare two groups. One-way or two-way ANOVA was used to compare multiple groups with post hoc Tukey test. *p* < 0.05 was considered significant. All data are shown as mean ± SEM and other details and the level of significance is indicated in figure legends.

## Study approval

The University of Pennsylvania institutional review board (IRB) approved the human kidney sample collection. We engaged with an external honest broker who was responsible for human kidney sample collection. No personal data were acquired.

## Data availability

GWAS, kidney QTL, snATAC-seq, and snRNA-seq data were published (GCST90100220, GSE115098, GSE173343, GSE172008 and GSE200547) earlier and are publicly available online at the Susztaklab.com Kidney Biobank. Additional details on protocols, and special reagents regarding this study are provided upon request to the corresponding author.

## Acknowledgments

This study was led by K.S.. K.S. and X.L. designed the study, analyzed the data, and wrote the manuscript with input from other authors. X.L., A.A., and A. N., performed experiments. H.H., H. L, J.Z., X.L., and K. A. performed computation analysis.

